# Automated Landmark-Based Root Inoculation in *Arabidopsis* Using Computer Vision and Robotics

**DOI:** 10.64898/2026.03.27.714898

**Authors:** Francisco Mansilha, Fedor Chursin, Borislav Nachev, Wesley van Gaalen, Vlad Matache, Vinicius Lube, Dean van Aswegen, Donald Jason Harty, Jason van Hamond, Valerian Meline, Marciel P. Mendes, M. Alican Noyan

## Abstract

Manual inoculation of plant roots is labor-intensive, spatially imprecise, and limits experimental throughput in plant–microbe interaction studies. Here, we present an integrated computer vision and robotics pipeline for automated, landmark-based root inoculation in *Arabidopsis thaliana*. Seedlings grown on Gelrite plates were imaged using the HADES automated phenotyping platform at the Netherlands Plant Eco-Phenotyping Centre, Utrecht University. A U-Net-based segmentation model (RootNet, F1 = 0.80) identified root structures, from which primary root tips were localized with a mean error of 0.25 mm. An affine coordinate transformation (mean target registration error: 1.09 mm) mapped image coordinates to the workspace of an Opentrons OT-2 liquid handling robot for targeted dispensing of 10 µL volumes. The system achieved successful inoculation in all 17 benchmark seedlings, corresponding to 100% accuracy (95% CI: 80–100%, Clopper-Pearson), and biological validation with fluorescent bacteria confirmed successful colonization along the root axis in 9 of 10 seedlings. To our knowledge, this is the first reported demonstration of automated, landmark-based root inoculation, extending the concept of automated phenotyping from passive measurement to active robotic intervention in real-time. The pipeline is generalizable to other root landmarks and organisms, enabling precise and reproducible delivery of microorganisms to specific root locations for systematic investigation of localized plant–microbe interactions.

## 1 Introduction

Plant pathogens and pests are estimated to cause an annual crop yield loss of $220 billion.^1^ With the added pressure of global warming and population growth, research on plant immunity and growth is becoming increasingly important for global food security. Plant-microbe interactions play an essential role in plant health.^2, 3^ Plants can use beneficial microbes to fight pathogens as well as to boost growth.^4, 5^ Moreover, researchers have long inoculated plants with pathogens to identify disease-resistant genotypes.^6, 7^ To facilitate this research, reliable delivery of beneficial and harmful microorganisms to defined plant tissues, a process known as inoculation, is necessary.

Inoculation is a technique widely used by plant scientists to study plant-microbe interactions. Organisms can be delivered before germination through seed inoculation, during growth through root or leaf inoculation, as well as through the growth medium such as soil inoculation.^8^ Each delivery route further splits into subcategories. For example, root inoculation can be done by dipping the whole root into the inoculant, known as root dip inoculation,^9^ or by delivering the inoculum to a specific location, referred to as spot or point inoculation.^10–12^ An optimal inoculation system must be rapid to facilitate high-throughput experiments such as large-scale screening of pathogen resistant phenotypes.^13, 14^ Additionally, it must ensure inoculum delivery of consistent doses to precise locations for reliable results across experiments.^4, 10^ Studies on inoculation methods for both beneficial and pathogenic organisms highlight that the delivery method affects the success of inoculation.^4, 13, 15–18^ Therefore, an optimal inoculation system must also allow testing multiple delivery methods and parameters.^19^ Given these requirements, there is a need for the development of automated inoculation systems.

When it comes to automation in plant science, the spotlight is on machine learning (ML) and robotics.^20, 21^ ML has been used to study molecules, individual plants, and crop fields, revolutionizing plant science across all scales.^22^ Robotic platforms have been developed for high-throughput root phenotyping. For example, RootBot automates the imaging of primary roots in drought-stress.^23^ However, these systems focus on imaging and measurement rather than physical intervention. In the biomedical domain, platforms such as LabDroid Maholo integrate robotics with artificial intelligence for automated cell culture,^24^ and RoboCulture combines a general-purpose robotic manipulator with computer vision for autonomous yeast culture experiments,^25^ demonstrating the feasibility of vision-guided liquid handling at millimeter precision. Combining plant localization and liquid delivery has been extensively demonstrated for crop and weed detection and selective spraying at the field scale.^26^

An automated inoculation system must be capable of accurately localizing an individual plant, identifying specific plant organs, and extracting the structural information of those organs. For root inoculation, accurate extraction of the root system architecture (RSA) is essential, and has been extensively studied in the literature.^27^ Lube et al. developed an image analysis software that can detect Arabidopsis seedlings in agar plates, find their shoot, seed and roots, and extract the root system architecture (RSA) hourly at scale.^28^ However, to our knowledge, no system has combined computer vision-based structural analysis of plant roots with targeted robotic inoculation at the individual-plant and organ level.

In this work, we propose an integrated computer vision and robotics pipeline (Figure 1) for the automated, landmark-based inoculation of plant roots. We define specific root landmarks, such as the primary root tip and junction, as explicit experimental parameters to guide precise liquid delivery. By automating these tasks, the system replaces intensive manual pipetting, reducing human labor while increasing the spatial consistency and reproducibility of the inoculation process. This system provides a foundation for experiments on localized inoculation, enabling precise delivery of microorganisms to user-defined root landmarks at defined developmental stages.

**Fig 1.**
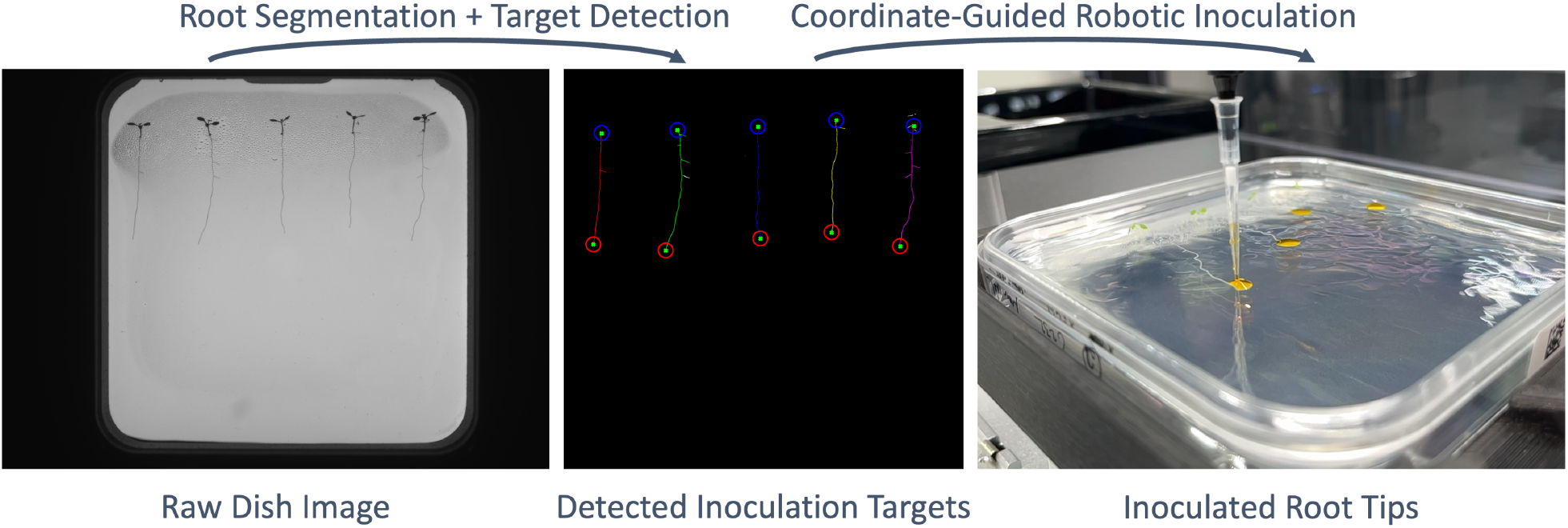
Overview of the integrated computer vision and robotics pipeline for automated landmark-based root inoculation. Arabidopsis seedlings grown on agar plates are imaged using the HADES platform. The computer vision pipeline performs root segmentation, skeletonization, and landmark extraction (primary root tip and junction). Image coordinates are transformed via an affine transformation to the Opentrons OT-2 robot workspace for targeted liquid delivery.

## 2 Materials and Methods

### 2.1 Datasets

We collected four datasets to support the development, evaluation, and validation of the proposed system (Table 1). All datasets were acquired using the HADES platform,^29^ a fully automated in vitro plant root phenotyping platform developed by Photon Systems Instruments (PSI) and operational at the Netherlands Plant Eco-phenotyping Centre (NPEC) located at Utrecht University. This facility is used for standardized studies of root development and plant–microbe interactions under sterile conditions.

**Table 1.** Summary of the datasets used in this study.

| Name | # Dish | # Seedling | Description |
| --- | --- | --- | --- |
| Development dataset | 428 | 1695 | Images with non-expert labeled root masks used for training and validation of the segmentation model |
| Expert benchmark dataset | 10 | 49 | Images with plant scientist annotations used to evaluate root length, tip, and junction prediction accuracy |
| Inoculation benchmark dataset | 4 | 17 | End-to-end system evaluation of vision-guided robotic inoculation using dye |
| Biological validation dataset | 2 | 10 | Qualitative assessment of bacterial growth following targeted root inoculation |

For the present study, HADES provided controlled cultivation, automated plate handling, and repeatable image acquisition of vertically grown Arabidopsis seedlings on agar plates, 5 seedlings per plate. Root morphology was recorded with a RootCam imaging module equipped with a 12.36 MP monochrome CMOS sensor, generating images at approximately 4202 × 3006 px under transmitted backlighting. In addition, fluorescence images were obtained using the same camera in combination with programmable excitation sources and selectable emission filters. The fluorescence images were used in the biological validation stage to confirm localized colonization after robotic application.

The *development dataset* was used for training and validation of the root segmentation model. It consists of 428 Petri dishes containing 1,695 seedlings, for which binary root masks were annotated by non-expert annotators. This dataset was split into training (342 images) and validation subsets (86 images) during model development to optimize segmentation performance, with the F1 score used as the primary evaluation metric.

The *expert benchmark dataset* consists of 10 Petri dishes (49 seedlings), annotated by plant scientists. In addition to segmentation masks, this dataset includes biologically relevant annotations such as primary root length, primary root tip location, and primary root-hypocotyl junction position. It was used to evaluate the accuracy of the computer vision pipeline in terms of biologically meaningful metrics, including root length error, primary root tip localization error, and junction localization error.

The *inoculation benchmark dataset* consists of 4 Petri dishes (17 seedlings after excluding non-viable samples) and was used for end-to-end evaluation of the integrated computer vision and robotics pipeline. For each dish, images acquired from HADES were processed to extract root tip locations, which were subsequently transformed into robot coordinates and used to guide an Opentrons OT-2 liquid handling system, a separate robotic platform external to HADES, for targeted inoculation with a dye.

Finally, the *biological validation dataset* includes 2 Petri dishes (10 seedlings) and was used to qualitatively assess the biological relevance of the system. These dishes were first imaged in grayscale, and the resulting images were used as input to the inoculation pipeline to guide bacterial application. After inoculation, the same dishes were re-imaged in HADES using fluorescence imaging, where higher fluorescence intensity indicated stronger localized bacterial presence along the root axis.

### 2.2 Image preprocessing

The raw grayscale images containing the Petri dishes had a resolution of 4202 × 3006 px (width × height). The dish sits at the center with a black region from the sample stage surrounding it. The Petri dish was cropped to remove the black region and reduce the computational load. The dish boundary was detected by analyzing intensity profiles along the horizontal and vertical axes and identifying transitions corresponding to the dish edges. The resulting bounding box is adjusted to enforce a square region. To enable patch-based processing, cropped images were padded such that both spatial dimensions are divisible by the patch size. This resulted in Petri dish images with a resolution of 2816 × 2816 px. Finally, the cropped and padded images were patched with a patch size of 256 × 256 px, yielding 121 patches per image.

### 2.3 Root segmentation

We developed a binary root segmentation model, called RootNet. It was implemented in Keras using TensorFlow as the backend. The model was based on a U-Net architecture^30^ with an encoder–decoder structure and skip connections between corresponding resolution levels. The encoder consisted of repeated blocks of two convolutional layers followed by max pooling, progressively increasing the number of feature channels from 16 to 256. The decoder mirrored this structure using transposed convolutions for upsampling, concatenation with encoder features, and convolutional refinement. Dropout layers were included after convolutional blocks to reduce over-fitting. The network takes grayscale images of size 256 × 256 as input and outputs a single-channel segmentation map of the same resolution. In total, the model contains 1,940,817 trainable parameters. A detailed layer-by-layer specification of the network architecture is provided in Supplementary Table S1. The model was trained using binary cross-entropy loss and the Adam optimizer with a learning rate of 10^−4^. A total of 41,382 training patches and 10,406 validation patches were used. Early stopping with a patience of 5 epochs was used.

### 2.4 Instance separation and structural extraction

The predicted segmentation masks were converted into individual plant instances using connected component analysis. Since the seeds were placed in fixed positions during dish preparation, seedling positions were consistent across Petri dishes. The images were partitioned into five equal regions along the horizontal axis, and a seedling ID was assigned to each region. The largest connected component within each region was assigned to the corresponding seedling ID. If a component was not connected to the root system, it was discarded.

To extract structural information, each root instance was skeletonized to obtain a one-pixel-wide representation. This skeleton was interpreted as a graph in which nodes correspond to pixels and edges represent connectivity. The highest node was predicted as the junction location and the lowest node was predicted as the primary root tip. The primary root was predicted as the shortest path between the junction and the primary root tip on this graph. The primary root length was calculated as the shortest path between these nodes, using Dijkstra’s algorithm implementation in NetworkX.^31^

### 2.5 Coordinate transformation and robotic inoculation

We used Opentrons® OT-2, a liquid handling robot, to automatically inoculate root tips. First, the image coordinates from the computer vision pipeline were converted to the robot workspace coordinates. This was followed by the OT-2 protocol we designed that can deliver liquids to given target coordinates.

To transform image coordinates to robot workspace coordinates, we prepared an empty Petri dish and manually marked eight calibration points. The calibration dish was imaged using HADES, and the image coordinates of each point were recorded. Coordinates were defined using the standard image coordinate system with origin at the top-left, and stored following array indexing notation as (*y*_*image*_, *x*_*image*_), corresponding to row and column indices.

The same calibration dish was then placed in the OT-2 workspace, and the pipette tip was manually positioned at each calibration point. The corresponding robot coordinates were recorded as (*x*_robot_, *y*_robot_) from the OT-2 coordinate system, where the origin is defined at the bottom-left of the deck coordinate frame, with the x-axis increasing to the right and the y-axis increasing away from the user. An affine transformation matrix was estimated using a least-squares approach, mapping image coordinates to robot coordinates. The transformation matrix was then fitted on all calibration data and used to convert target coordinates (*y*_*image*_, *x*_*image*_) in pixels from image space to robot workspace coordinates (*x*_robot_, *y*_robot_) in millimeters. Transformation accuracy was evaluated separately using 4-fold cross-validation, as described in the Evaluation section.

The transformed coordinates were subsequently used to guide a sterile Opentrons OT-2 liquid handling unit for targeted inoculation. We designed an inoculation protocol using the Opentrons Python Protocol API.^32^ In each cycle, the robot picked up a fresh 2-20 µL pipette tip, aspirated 10 µL of liquid from a source well in a 96-well plate, moved to the target coordinate, dispensed the liquid at the desired root location, and discarded the tip before proceeding to the next point. This process was repeated for all target points.

For inoculation benchmarking experiments, a lab-grade chemical dye (Orange G, Sigma-Aldrich) was used. The dye enabled clear visualization of deposited droplets in RGB images, to help accurate counting of successful inoculations. For biological validation, a bacterial inoculum, described in the next section, was used to test whether targeted robotic application could generate localized biological activity at the intended root position.

### 2.6 Bacterial inoculum preparation

Bacteria (*Pseudomonas simiae* WCS417 tagged with mCherry) were streaked from a glycerol stock onto Petri dishes containing agar-solidified King’s B medium supplemented with 50 *µ*g mL^−1^ rifampicin and incubated upside down overnight at 28 °C in darkness.

In a sterile hood, 2–5 mL of sterile 10 mM MgSO_4_ was added to the overnight culture, and bacteria were suspended by scraping them off the agar using a sterile spreader. A volume of 50 *µ*L of this suspension was plated onto a fresh KB plate containing rifampicin and spread evenly. Plates were incubated upside down overnight at 28 °C in darkness.

After 24 h, 5 mL of 10 mM MgSO_4_ was added, and bacteria were gently scraped into suspension. The suspension was transferred to a 50 mL tube and centrifuged at 4,500 *g* for 5 min. The supernatant was discarded, and the pellet was resuspended in 25 mL of 10 mM MgSO_4_ and centrifuged again. The pellet was then resuspended in 20 mL of fresh 10 mM MgSO_4_.

Optical density at 600 nm (OD_600_) was measured, and the suspension was diluted to OD_600_ = 0.1. The suspension was subsequently distributed into a 96-well plate.

### 2.7 Evaluation

We evaluated the end-to-end system performance as well as the performance of the subsystems. To evaluate the segmentation performance of RootNet, we calculated the F1 score on the *validation dataset*, which was a subset of the *development dataset*. To evaluate biologically relevant trait extraction, we measured primary root length absolute percentage error, localization errors for the junction and primary root tip (mm) using the *expert benchmark dataset*.

To evaluate the image-to-robot coordinate transformation, we used the calibration Petri dish described in the previous section. A 4-fold cross-validation procedure was used for evaluation. In each fold, six points were used to estimate the affine transformation, and the remaining two points were used for evaluation. The transformation accuracy was quantified using the target registration error (TRE), defined as the Euclidean distance between the predicted robot coordinates and the ground-truth robot coordinates of the held-out points. The TRE was computed for each fold and averaged across all folds to obtain the final transformation error.

The end-to-end system performance was evaluated by inoculating 17 seedlings at primary root tips and counting how many droplets covered the root tips. A lab-grade dye and an RGB mirrorless Canon R5 camera with an EF f/1.8 50mm lens were used to enable clear visualization of deposited droplets and accurate assessment of success right after inoculation. Accuracy was reported as the number of successful inoculations divided by the number of seedlings, using the *inoculation benchmark dataset*. Finally, a bacterial inoculum was delivered to the root tips of 10 seedlings, and fluorescence imaging was performed to qualitatively assess localized bacterial growth, using the *biological validation dataset*.

## 3 Results

The binary root segmentation model, RootNet, achieved an F1 score of 0.8042 and a binary cross-entropy loss of 0.0048 on the validation dataset. The *validation dataset* contained 10,406 patches from 86 images annotated by non-experts, which may introduce annotation noise. Therefore, we evaluated the biological trait extraction performance using *expert benchmark dataset*, which contained 49 seedlings labeled by experts.

On the expert benchmark dataset, the mean absolute percentage error (MAPE) for the length of the primary root was 2.90%. The localization error for the junction between the hypocotyl and the primary root was 0.66 mm. The mean primary root tip localization error was 0.25 mm. The primary root tip localization error distribution included a single large outlier (max: 4.75 mm), while the remaining samples showed low localization error. Figure 2 shows the output of the computer vision pipeline on 3 of the 10 images in the expert benchmark dataset. Red circles indicate the predicted primary root tip location and the blue circles indicate the predicted junction location. Table 2 shows the biologically relevant trait extraction results, including the mean, median, P10, and P90.

**Table 2.** Expert Benchmark Results.

| Metric | Mean | Median | P10 | P90 |
| --- | --- | --- | --- | --- |
| Primary root length absolute percentage error (%) | 2.90 | 1.23 | 0.24 | 8.33 |
| Junction localization error (mm) | 0.66 | 0.37 | 0.06 | 1.66 |
| Primary root tip localization error (mm) | 0.25 | 0.12 | 0.04 | 0.22 |

**Fig 2.**
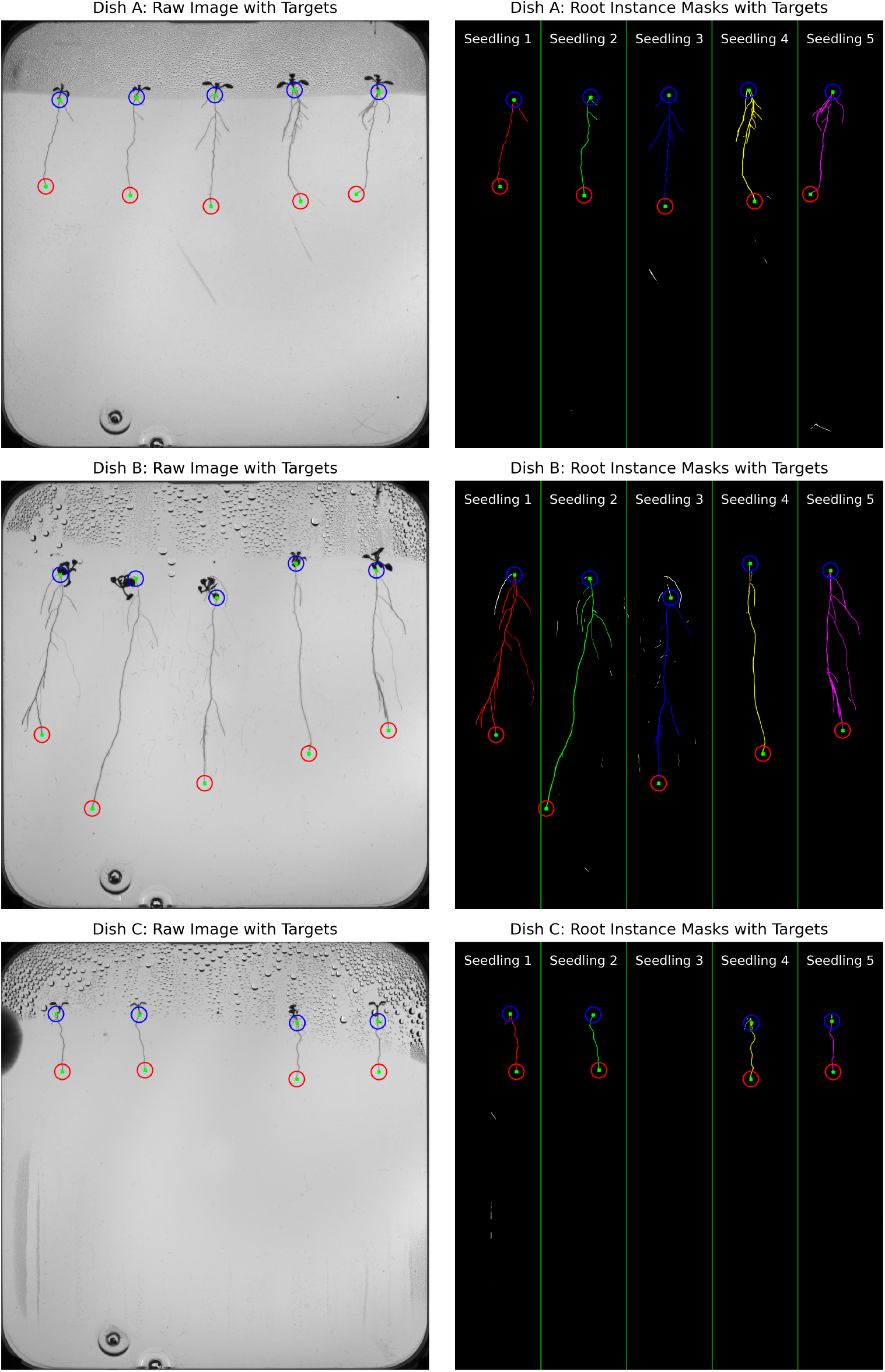
Output of the computer vision pipeline on three selected dishes from the expert benchmark dataset. Red circles indicate predicted primary root tip locations; blue circles indicate predicted junction locations. In Dish B, the top-left lateral root of Seedling 1 (white) is disconnected from the root system (red) due to shoot occlusion. For Seedling 3 in Dish B, the shoot folded over the primary root, displacing the predicted junction downward.

We used 8 calibration points to quantify the performance of the image-to-robot coordinate transformation. The resulting target registration error was 1.09 mm on average, with a minimum of 0.75 mm and a maximum of 1.43 mm. This error reflects the combined effect of transformation estimation, imaging resolution, and robotic positioning accuracy, and defines a practical lower bound for the transformation component of the full system.

The end-to-end system performance was evaluated using the *inoculation benchmark dataset* and inoculating 17 seedlings with an orange dye. The system took HADES images as input, predicted the root tip coordinates, and provided these coordinates to the OT-2 inoculation protocol, resulting in targeted inoculation of root tips. After inoculation, RGB images of the plates were acquired and the number of successful inoculations was counted. An inoculation was considered successful if the root tip was within the droplet. We observed 17 successful inoculations out of 17 seedlings, corresponding to 100% accuracy. Figure 3 shows an example plate from the *inoculation benchmark dataset*, with a close-up illustrating a successful inoculation.

**Fig 3.**
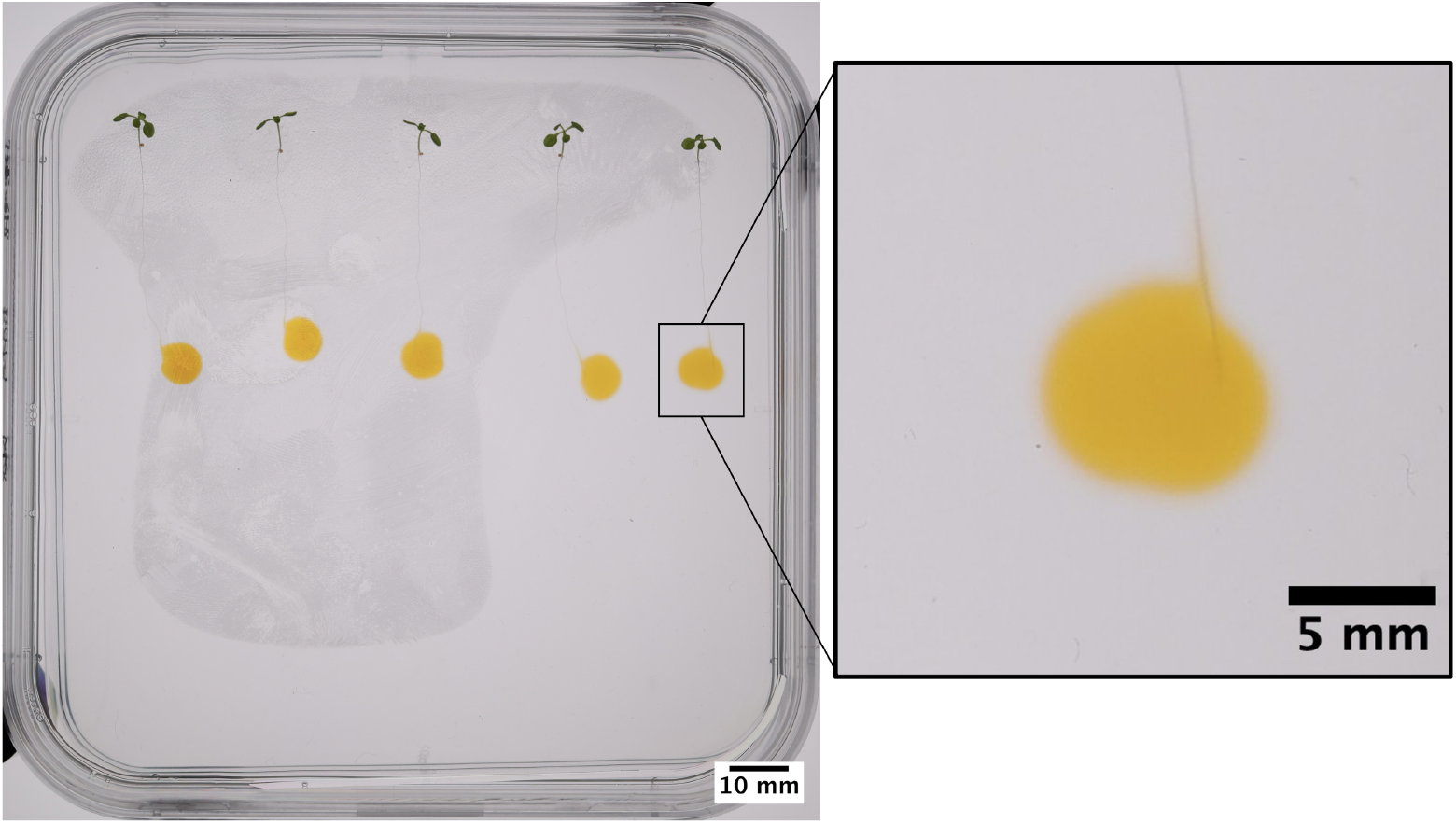
Dye inoculation results from the inoculation benchmark dataset. (a) Overview of a Petri dish after targeted inoculation with Orange G dye at predicted root tip locations. (b) Close-up view showing a successful inoculation where the 10 µL dye droplet covers the primary root tip. All 17 seedlings across 4 dishes were successfully inoculated.

Finally, the 10 seedlings in the *biological validation dataset* were inoculated automatically with the described system to assess bacterial growth. A fluorescence signal was observed along the primary root axis for 9 of the seedlings out of 10. One seedling was contaminated during the experiment. Figure 4 shows both the grayscale and fluorescence images of one of the dishes. Among the five seedlings in the imaged dish, four displayed clear fluorescence signals indicative of successful colonization. The contaminated seedling can also be seen in Figure 4a.

**Fig 4.**
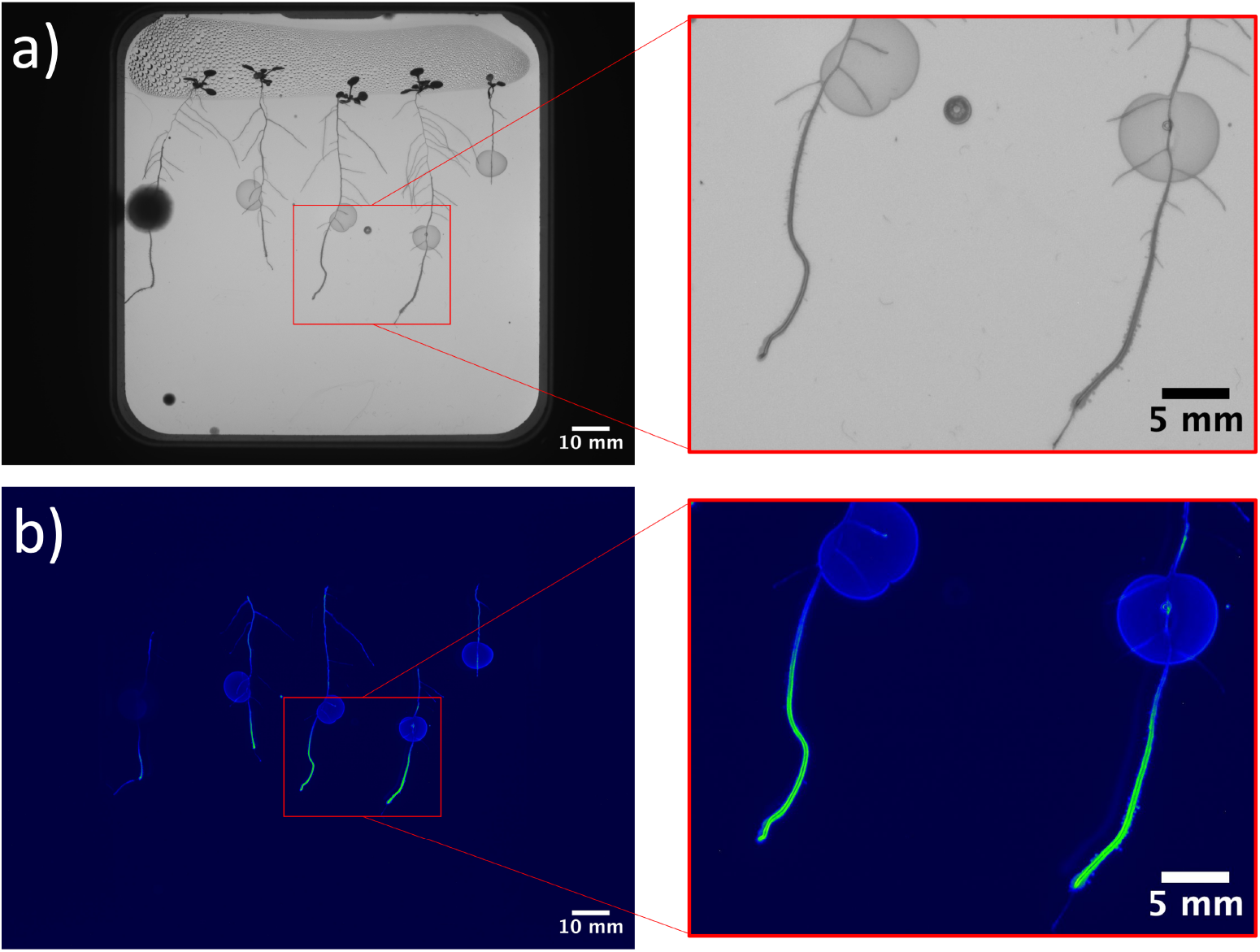
Biological validation of targeted root inoculation. (a) Grayscale image of a Petri dish with five Arabidopsis seedlings prior to inoculation. The leftmost seedling on the dish was contaminated. (b) Fluorescence image of the same dish after bacterial inoculation, showing fluorescent signal along the primary root axis of four seedlings, indicating successful bacterial colonization at the targeted locations.

## 4 Discussion

This study demonstrates an integrated computer vision and robotics pipeline for targeted, landmark-based root inoculation on agar plates. The results show that primary root tips can be localized with high accuracy and inoculated reliably, supporting the feasibility of automated and higher-throughput localized inoculation experiments.

The overall performance of the system depends on multiple sequential components: image quality, segmentation performance, root system architecture extraction, coordinate transformation, and the inoculation protocol. Errors in earlier stages propagate through the pipeline, making robustness at each step important for end-to-end performance.

Image quality has a direct effect on segmentation. Artifacts such as scratches, bubbles, impurities like paper fibers, condensation, or contamination can reduce segmentation accuracy. In addition, discontinuities in the segmentation mask lead to incomplete root representations. These discontinuities are often caused by occlusions, for example when the shoot detaches from the agar and folds over the root system. This results in missing connections in the root mask and affects downstream processing. As shown in Figure 2, the top-left lateral root of Dish B Seedling 1 (white) is not connected to the root system (red) due to shoot occluding the connection. For Seedling 3 in Dish B, the shoot folded over to the primary root, causing the predicted junction point to be much lower than its actual position. This limitation contributes to higher errors in junction detection compared to root tip detection. The junction localization error (0.66 mm) is 2.6 times higher than the root tip error (0.25 mm). The pipeline currently lacks a mechanism to restore such discontinuities, which limits performance in these scenarios.

The primary root tip localization metric contained a single large outlier (4.75 mm), corresponding to Seedling 1 in Figure 2 (Dish B). In this case, the primary root grew at a slight angle, while a lateral root extended further downward in the gravity direction. Because the pipeline defines the lowest graph node as the primary root tip, it selected the lateral root tip instead of the true primary root tip. This reflects a limitation of the current rule-based landmark definition rather than a failure to detect a root tip.

The transformation from image to robot coordinates introduces an additional source of error, with a mean target registration error of 1.09 mm. This represents a practical lower bound on inoculation precision and reflects combined uncertainties from calibration, imaging resolution, and robotic positioning.

The perfect success rate (17/17), while encouraging, likely reflects the favorable ratio between droplet size and spatial error. A 10 µL droplet on agar spreads to roughly 5 mm in diameter, providing a tolerance margin relative to the combined localization and target registration error. Under more demanding conditions, such as smaller volumes or more closely spaced landmarks, the current error budget would likely become limiting. As a proof-of-concept demonstration, 100% success on 17 seedlings (95% CI: 80–100%, Clopper-Pearson) is promising, but larger datasets will be needed for statistically tighter performance estimates.

The instance segmentation approach assumes a fixed spatial arrangement of seedlings and relies on selecting the largest connected component per region. This assumption restricts applicability in cases where seedlings overlap or deviate from expected positions. Overlapping root systems were excluded from evaluation, and extending the method to handle such cases remains an important direction for future work.

The current benchmark system required manual transfer of plates between HADES and the OT-2, which limits full autonomy and throughput. However, HADES already includes robotic plate-handling infrastructure, which could support future integration of imaging, analysis, and inoculation into a fully automated workflow without human intervention. This integration is currently being explored at NPEC.

Biological validation confirmed that targeted inoculation leads to successful bacterial colonization along the root axis, as indicated by fluorescence imaging. However, this validation was qualitative and performed on a limited number of samples (9 of 10 seedlings showed colonization). Future work should include quantitative assessment of bacterial load, spatial distribution, and downstream plant responses across larger datasets to fully establish biological reliability.

The development dataset was annotated by non-expert annotators, which may introduce annotation noise, including missing thin or short lateral roots, inconsistent segmentation at root tips, and inaccurate placement of the hypocotyl–root junction. However, the expert benchmark dataset serves as an independent quality check, and the low root tip localization error (0.25 mm) suggests that segmentation quality is sufficient for the downstream inoculation task. Future work should report inter-annotator agreement to quantify annotation reliability.

Finally, while this study focuses on root tip inoculation, the pipeline is generalizable to other landmarks within the root system. This enables systematic investigation of how inoculation location influences plant–microbe interactions, which is difficult to achieve with manual methods. Extending the framework to support multiple target locations and experimental conditions will further enhance its utility for plant science research.

## 5 Conclusion

This study presented an integrated computer vision and robotics pipeline for automated, landmark-based inoculation of *Arabidopsis* root tips on agar plates. The system achieved sub-millimeter root tip localization accuracy and enabled successful inoculation across all benchmark samples. Biological validation confirmed successful bacterial colonization at targeted root locations via fluorescence imaging.

To our knowledge, this is the first reported demonstration of automated, landmark-based root inoculation, extending automated phenotyping from passive measurement to active robotic intervention. By linking computer vision-based root analysis to precise physical inoculation, the system enables scalable, reproducible delivery of microorganisms to specific root locations.

The proposed framework is generalizable to other root landmarks, plant species, and experimental conditions, providing a foundation for systematic investigation of localized plant–microbe interactions.

## 6 Code, Data, and Materials Availability

Code and data supporting this study are available upon reasonable request.

## 7 Acknowledgments

We thank the Netherlands Plant Eco-phenotyping Centre (NPEC) at Utrecht University for providing access to the HADES platform and for supporting experimental coordination. We also thank colleagues from the Plant–Microbe Interactions group at Utrecht University for assistance with bacterial strains and fluorescence-imaging workflows. This work was supported by the AI Experiments Fund at Breda University of Applied Sciences.

## 8 Author Contributions

F.M., F.C., B.N., W.v.G., V. Matache, and M.A.N. developed the computer vision and robotics pipeline. D.v.A. and D.J.H. contributed to the simulation of the Opentrons OT-2 environment and supported validation of the inoculation pipeline prior to physical deployment. J.v.H. contributed to methodology development and experimental validation. V. Meline provided biological inputs, including protocols for fluorescence data extraction. M.P.M. contributed biological supervision, resources, and operation of the HADES platform. V.L. contributed conceptualization of the vision-guided robotic inoculation framework, access to HADES datasets, project administration, and infrastructure support from NPEC. M.A.N. contributed conceptualization, supervision, project administration, and writing, review, and editing. All authors reviewed and approved the final manuscript.

