## Supplementary material for "Automated Landmark-Based Root Inoculation in *Arabidopsis* Using Computer Vision and Robotics": Table S1

### Supplementary Materials

**Table S1** Model architecture.

| Layer (type) | Output Shape | Param # | Connected to |
|---|---|---|---|
| input_1 (InputLayer) | (None, 256, 256, 1) | 0 | – |
| conv2d (Conv2D) | (None, 256, 256, 16) | 160 | input_1 |
| dropout (Dropout) | (None, 256, 256, 16) | 0 | conv2d |
| conv2d_1 (Conv2D) | (None, 256, 256, 16) | 2320 | dropout |
| max_pooling2d (MaxPooling2D) | (None, 128, 128, 16) | 0 | conv2d_1 |
| conv2d_2 (Conv2D) | (None, 128, 128, 32) | 4640 | max_pooling2d |
| dropout_1 (Dropout) | (None, 128, 128, 32) | 0 | conv2d_2 |
| conv2d_3 (Conv2D) | (None, 128, 128, 32) | 9248 | dropout_1 |
| max_pooling2d_1 (MaxPooling2D) | (None, 64, 64, 32) | 0 | conv2d_3 |
| conv2d_4 (Conv2D) | (None, 64, 64, 64) | 18496 | max_pooling2d_1 |
| dropout_2 (Dropout) | (None, 64, 64, 64) | 0 | conv2d_4 |
| conv2d_5 (Conv2D) | (None, 64, 64, 64) | 36928 | dropout_2 |
| max_pooling2d_2 (MaxPooling2D) | (None, 32, 32, 64) | 0 | conv2d_5 |
| conv2d_6 (Conv2D) | (None, 32, 32, 128) | 73856 | max_pooling2d_2 |
| dropout_3 (Dropout) | (None, 32, 32, 128) | 0 | conv2d_6 |
| conv2d_7 (Conv2D) | (None, 32, 32, 128) | 147584 | dropout_3 |
| max_pooling2d_3 (MaxPooling2D) | (None, 16, 16, 128) | 0 | conv2d_7 |
| conv2d_8 (Conv2D) | (None, 16, 16, 256) | 295168 | max_pooling2d_3 |
| dropout_4 (Dropout) | (None, 16, 16, 256) | 0 | conv2d_8 |
| conv2d_9 (Conv2D) | (None, 16, 16, 256) | 590080 | dropout_4 |
| conv2d_transpose (Conv2DTranspose) | (None, 32, 32, 128) | 131200 | conv2d_9 |
| concatenate (Concatenate) | (None, 32, 32, 256) | 0 | conv2d_transpose, conv2d_7 |
| conv2d_10 (Conv2D) | (None, 32, 32, 128) | 295040 | concatenate |
| dropout_5 (Dropout) | (None, 32, 32, 128) | 0 | conv2d_10 |
| conv2d_11 (Conv2D) | (None, 32, 32, 128) | 147584 | dropout_5 |
| conv2d_transpose_1 (Conv2DTranspose) | (None, 64, 64, 64) | 32832 | conv2d_11 |
| concatenate_1 (Concatenate) | (None, 64, 64, 128) | 0 | conv2d_transpose_1, conv2d_5 |
| conv2d_12 (Conv2D) | (None, 64, 64, 64) | 73792 | concatenate_1 |
| dropout_6 (Dropout) | (None, 64, 64, 64) | 0 | conv2d_12 |
| conv2d_13 (Conv2D) | (None, 64, 64, 64) | 36928 | dropout_6 |
| conv2d_transpose_2 (Conv2DTranspose) | (None, 128, 128, 32) | 8224 | conv2d_13 |
| concatenate_2 (Concatenate) | (None, 128, 128, 64) | 0 | conv2d_transpose_2, conv2d_3 |
| conv2d_14 (Conv2D) | (None, 128, 128, 32) | 18464 | concatenate_2 |
| dropout_7 (Dropout) | (None, 128, 128, 32) | 0 | conv2d_14 |
| conv2d_15 (Conv2D) | (None, 128, 128, 32) | 9248 | dropout_7 |
| conv2d_transpose_3 (Conv2DTranspose) | (None, 256, 256, 16) | 2064 | conv2d_15 |
| concatenate_3 (Concatenate) | (None, 256, 256, 32) | 0 | conv2d_transpose_3, conv2d_1 |
| conv2d_16 (Conv2D) | (None, 256, 256, 16) | 4624 | concatenate_3 |
| dropout_8 (Dropout) | (None, 256, 256, 16) | 0 | conv2d_16 |
| conv2d_17 (Conv2D) | (None, 256, 256, 16) | 2320 | dropout_8 |
| conv2d_18 (Conv2D) | (None, 256, 256, 1) | 17 | conv2d_17 |
| **Total params: 1,940,817 (7.40 MB)**<br>**Trainable params: 1,940,817 (7.40 MB)**<br>**Non-trainable params: 0 (0.00 Byte)** | | | |
